# Multiple event segmentation mechanisms in the human brain

**DOI:** 10.1101/2025.07.07.663487

**Authors:** Tan T. Nguyen, Joset A. Etzel, Matthew A. Bezdek, Jeffrey M. Zacks

## Abstract

The human brain segments continuous experience into discrete events, with theoretical accounts proposing two distinct mechanisms: creating boundaries at points of high *prediction error* (mismatch between expected and observed information) or high *prediction uncertainty* (reduced precision in predictions). Using fMRI and computational modeling, we investigated the neural correlates of error-driven and uncertainty-driven boundaries. We developed computational models that generate boundaries based on prediction error or prediction uncertainty, and examined how both types of boundaries, and human-identified boundaries, related to fMRI pattern shifts and evoked responses. Multivariate analysis revealed a specific temporal sequence of neural pattern changes around human boundaries: early pattern shifts in anterior temporal regions (-11.9s), followed by shifts in parietal areas (-4.5s), and subsequent whole-brain pattern stabilization (+11.8s). The core of this dynamic response was associated with both error-driven and uncertainty-driven boundaries. Critically, both error- and uncertainty-driven boundaries were associated with unique pattern shifts. Error-driven boundaries were associated with early pattern shifts in ventrolateral prefrontal areas, followed by pattern stabilization in prefrontal and temporal areas. Uncertainty-driven boundaries were linked to shifts in parietal regions within the dorsal attention network, with minimal subsequent stabilization. In addition, within the core regions responsive to both types of boundaries, the timing differed significantly. These findings provide evidence for two overlapping brain networks that maintain and update representations of the environment, controlled by two distinct prediction quality signals: prediction error and prediction uncertainty.

## Introduction

To act effectively in dynamic environments, humans use memory and knowledge to predict how activity will unfold over time (Clark, 2013; Friston et al., 2016; Knott & Takac, 2021; Niv & Schoenbaum, 2008). Contemporary cognitive models propose that perception is guided by event models—stable representations of the current situation (Zacks, 2020). These event models integrate sensory information with prior schematic knowledge—structured representations that encode knowledge about event classes (Anderson, 1978; Bartlett, 1932; Graesser & Nakamura, 1982). For example, when serving coffee, an event model helps predict that the recipient will pick up the cup and how they will move to do so. However, for such models to remain effective, they must update appropriately when one event ends and another begins—once the coffee has been taken, a coffee-serving model becomes unhelpful (DuBrow et al., 2017); this is known as *event segmentation*. There has been considerable recent interest in the features that predict event segmentation (Bailey & Zacks, 2015; Brunec et al., 2018; Magliano et al., 1999; Su & Swallow, 2024; Wang & Egner, 2022), in the effects of segmentation on immediate access to information in working memory (Radvansky et al., 2010; Radvansky & Copeland, 2006), and in the role of segmentation in shaping long-term memory (Clewett et al., 2019; Ding & Zacks, 2025; DuBrow & Davachi, 2013; Sargent et al., 2013; Swallow et al., 2009). Individual and group differences in segmentation are robust predictors of memory and action performance. However, an important open question is: What control signal or signals does the cognitive system use to determine when to update event models?

One potential signal to control event model updating is *prediction error*—the difference between the system’s prediction and what actually occurs. A transient increase in prediction error is a valid indicator that the current model no longer adequately captures the current activity. Event Segmentation Theory (EST; Zacks et al., 2007) proposes that event models are updated when prediction error increases beyond a threshold, indicating that the current model no longer adequately captures ongoing activity. A related but computationally distinct proposal is that prediction *uncertainty* (also termed “unpredictability”) can serve as a control signal (Baldwin & Kosie, 2021). The advantage of relying on prediction uncertainty to detect event boundaries is that it is inherently proactive: the cognitive system can start looking for cues about what might come next before the next event starts (Baldwin & Kosie, 2021; Kuperberg, 2021). Building on this idea, Gumbsch et al. (2022) developed a computational model in which uncertainty dynamically directs the model’s gaze, which successfully predicted infant gaze allocation.

In naturalistic activity, prediction error and prediction uncertainty tend to co-occur. In a recent study (Nguyen et al., 2024), we used a large corpus and a formal computational model to assay the joint potential contributions of prediction error and prediction uncertainty to human event segmentation. We found that both error-driven and uncertainty-driven updating signals aligned with human segmentation and categorization judgments, but that uncertainty-driven segmentation aligned significantly better. This result argues for a unique role for prediction uncertainty in event model updating. However, it also opens up the possibility that multiple control mechanisms could govern updating of stable brain representations. One possibility is that representational updates in some brain areas may be triggered by either prediction error spikes or prediction uncertainty spikes. Another possibility (not mutually exclusive) is that updates in some brain areas may be uniquely triggered by prediction error or prediction uncertainty.

A body of fMRI research has shown that event boundaries evoke responses in multiple brain regions. Early univariate analyses found transient increases in BOLD activity at event boundaries within visual and parietal areas, including medial visual cortex, the precuneus, and intraparietal sulcus (Speer et al., 2007; Zacks, 2010; Zacks et al., 2001). Moreover, regions within the default mode network showed decreased activity at musical and movie event boundaries (Burunat et al., 2024; Zacks, 2010). Boundary-related activity is predictive of subsequent memory, supporting a causal role for segmentation in memory formation (Ben-Yakov & Dudai, 2011). Recent research has further revealed that hippocampal responses at movie event boundaries show a graded profile, wherein greater univariate activity is associated with greater perceived boundary “salience,” measured by agreement across a separate group of individuals that a boundary occurred (Ben-Yakov & Henson, 2018).

More recently, multivariate approaches have provided insights into neural representations during event segmentation. One prominent approach uses hidden Markov models (HMMs) to detect moments when the brain switches from one stable activity pattern to another (Baldassano et al., 2017) during movie viewing; these periods of relative stability were referred to as “neural states” to distinguish them from subjectively perceived events. Sensory regions like visual and auditory cortex showed faster transitions between neural states. Multi-modal regions like the posterior medial cortex, angular gyrus, and intraparietal sulcus showed slower neural state shifts, and these shifts aligned with subjectively reported event boundaries. Geerligs et al. (2021, 2022) employed a different analytical approach called Greedy State Boundary Search (GSBS) to identify neural state boundaries. Their findings echoed the HMM results: short-lived neural states were observed in early sensory areas (visual, auditory, and somatosensory cortex), while longer-lasting states appeared in multi-modal regions, including the angular gyrus, posterior middle/inferior temporal cortex, precuneus, anterior temporal pole, and anterior insula. Particularly prolonged states were found in higher-order regions such as lateral and medial prefrontal cortex. They found that transitions between neural states in the default mode network (DMN) significantly coincide with human-identified event boundaries. Geerligs et al. (2022) further mapped the specific brain regions showing statistically significant overlap between neural state boundaries and perceived event boundaries, identifying the anterior cingulate cortex, dorsal medial prefrontal cortex, left superior and middle frontal gyrus, and anterior insula as areas with particularly strong alignment. These findings suggest that the subjective experience of event boundaries may be supported by shifts in neural activity patterns in these brain regions.

The previous evidence about evoked responses at event boundaries indicates that these are dynamic phenomena evolving over many seconds, with different brain areas showing different dynamics (Ben-Yakov & Henson, 2018; Burunat et al., 2024; Kurby & Zacks, 2018; Speer et al., 2007; Zacks, 2010). Less is known about the dynamics of pattern shifts at event boundaries (e.g. whether shifts observed in higher-order regions precedes or follow shifts observed in lower-level regions), because the HMM and GSBS analysis methods do not directly provide moment-by-moment measures of pattern shifts. Both the spatial and temporal aspects of evoked responses and pattern shifts at event boundaries have the potential to provide evidence about two potential control processes (error-driven and uncertainty-driven) for event model updating. However, this requires an approach that can model each potential mechanism independently and predict when event model updating driven by each mechanism should happen. Here, we combined fMRI with quantitative computational models of event segmentation that had been previously validated against human behavior (Nguyen et al., 2024). The models make predictions about when error-driven updating or uncertainty-driven updating should lead to fMRI evoked responses and pattern shifts. This approach allowed us to characterize the spatiotemporal profiles of neural dynamics associated with event segmentation.

Our study addresses three main research questions. First, how do neural activity and neural activity pattern change before, during, and after human-identified event boundaries? We aimed to replicate and extend previous findings using both univariate and multivariate analyses. Second, are human-identified boundaries predicted by both error- and uncertainty-driven computational models? We compared the alignment between our model-derived boundaries and the segmentation judgments of human observers, testing whether both error and uncertainty signals contribute uniquely to subjective experience of event boundaries. Third, and most importantly, do error- and uncertainty-driven boundaries engage distinct brain networks with different temporal dynamics? By comparing brain regions whose neural pattern changes are related to error-driven or uncertainty-driven boundaries and the temporal profiles of neural pattern changes, we can determine whether these computational processes rely on separate neural systems with distinct temporal signatures.

## Results

In this study, we examined brain responses around event boundaries during naturalistic video viewing. Forty-five participants watched four videos depicting everyday activities while undergoing fMRI scanning (see Methods). For each activity, a separate group of 30 participants had previously segmented each movie to identify fine-grained event boundaries (Bezdek et al., 2022). The mean event length was 21.4 s (median 22.2 s, SD 16.1 s). Mean event lengths for uncertainty-driven model and error-driven model were 28.96s, and 24.7s, respectively (Nguyen et al., 2024). We applied Finite Impulse Response (FIR) models to analyze brain activity changes and neural pattern shifts within a window of approximately 20 seconds before and after human boundaries, error-driven boundaries, and uncertainty-driven boundaries. For all analyses, we described activity within regions defined by the Schaefer 400-parcel atlas (Schaefer et al., 2018).

Our analyses below addressed three main questions about the neural mechanisms of event segmentation. We first characterized how the magnitude and pattern of neural activity change around human-identified event boundaries. Next, we tested whether human event segmentation might be supported by one or both candidate cognitive mechanisms by examining the extent to which error-driven and uncertainty-driven boundaries could predict human boundaries. Finally, we investigated whether these two potential control mechanisms engage separate neural systems by comparing the brain networks and temporal dynamics associated with error-driven and uncertainty-driven boundaries.

### Event boundaries were associated with increased activity in regions within the visual and dorsal attention networks, and decreased activity in regions within the default network

We first analyzed the BOLD response around human-identified event boundaries. We analyzed two effects: changes in BOLD magnitude (this section) and in local patterns of activity (following section). The magnitude analyses are a replication of previous studies (Burunat et al., 2024; Kurby & Zacks, 2018; Speer et al., 2007; Zacks, 2010): We time-locked the magnitude of the evoked response to the identified boundaries. Whereas previous studies were analyzed at the voxel level, here we took advantage of modern functional anatomic parcellation (Schaefer et al., 2018) to improve sensitivity. The pattern-based analyses represent a significant new view of the brain’s response to event boundaries. In the next section, we time-locked the rate of pattern change to the event boundary. This constitutes a valuable complement to methods based on the HMM (Baldassano et al., 2017) or GSBS (Geerligs et al., 2021) by providing a continuous time course of pattern change around the boundary.

To analyze changes in BOLD magnitude, we employed a finite impulse response (FIR) model over a time window spanning approximately 20 seconds before and 20 seconds after human-identified event boundaries (see Methods). This analysis revealed brain regions whose BOLD activity changes significantly (FDR with alpha < 0.001, corrected-p-value < 0.001) around event boundaries. The strongest changes were in areas within the visual and dorsal attention networks; the precuneus and medial prefrontal cortex (PFC) within the default network, and lateral PFC within the control network also had significant activity changes (Figure 1). The spatial distribution of these responses are similar to previous results (Burunat et al., 2024; Kurby & Zacks, 2018; Speer et al., 2007; Zacks, 2010).

**Figure 1:**
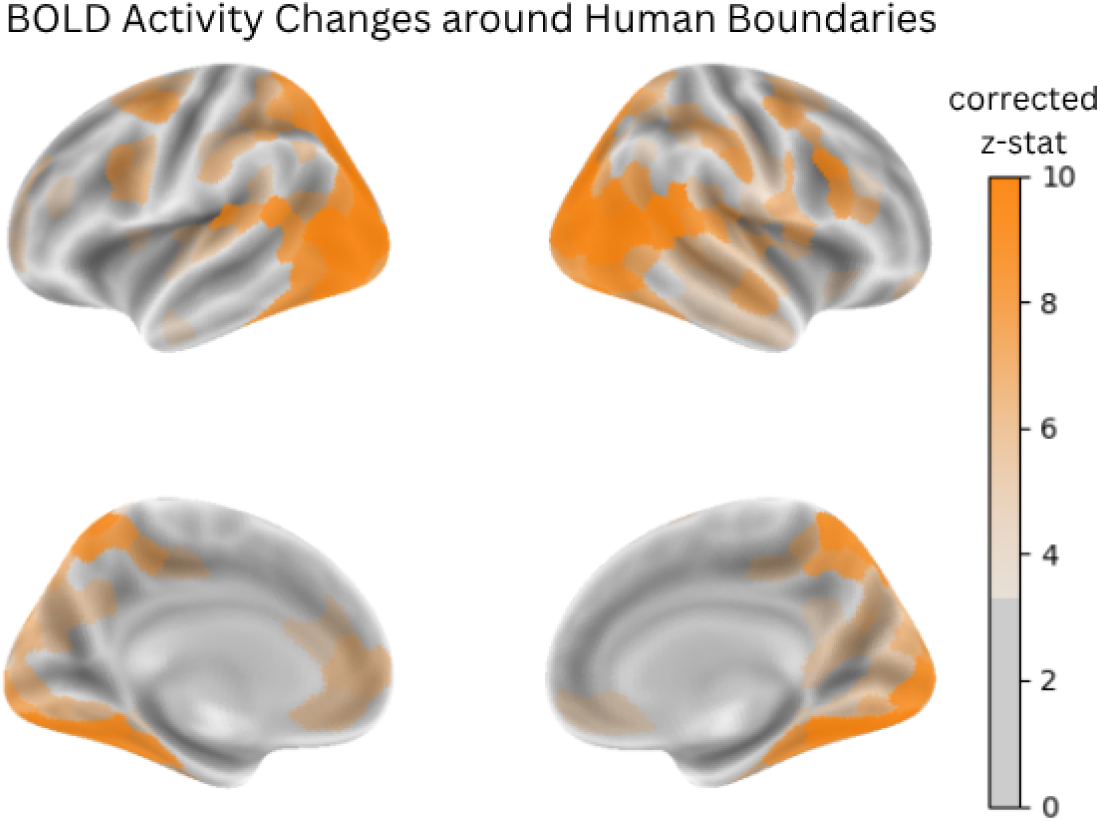
Regions where BOLD activity changed significantly (corrected p < 0.001) around human boundaries. FIR models were used to predict parcel-wise BOLD activity within a time window spanning approximately 20 seconds before and 20 seconds after human-identified event boundaries. The z-statistics revealed significant activity changes across multiple brain networks, with particularly strong changes in the visual and dorsal attention networks, and moderate changes in the precuneus and medial PFC within the default model network, and lateral PFC within the control network.

We then examined these evoked responses in detail, extracting FIR-estimated BOLD activity timecourses for each parcel showing significant change over time. Parcels in the visual, dorsal, and control networks showed an increase in BOLD activity around event boundaries, whereas many default network parcels showed decreases in BOLD activity around event boundaries (Figure 2). Interestingly, some regions, such as the precuneus, displayed a biphasic response characterized by an initial increase in activity at the event boundary followed by a subsequent decrease (Figure 2, bottom). Again, these findings are consistent with prior studies examining brain activity at event boundaries (Burunat et al., 2024; Kurby & Zacks, 2018; Speer et al., 2007; Zacks, 2010), and suggest that event segmentation is a process that unfolds over more than 20 seconds, involving multiple brain networks.

**Figure 2:**
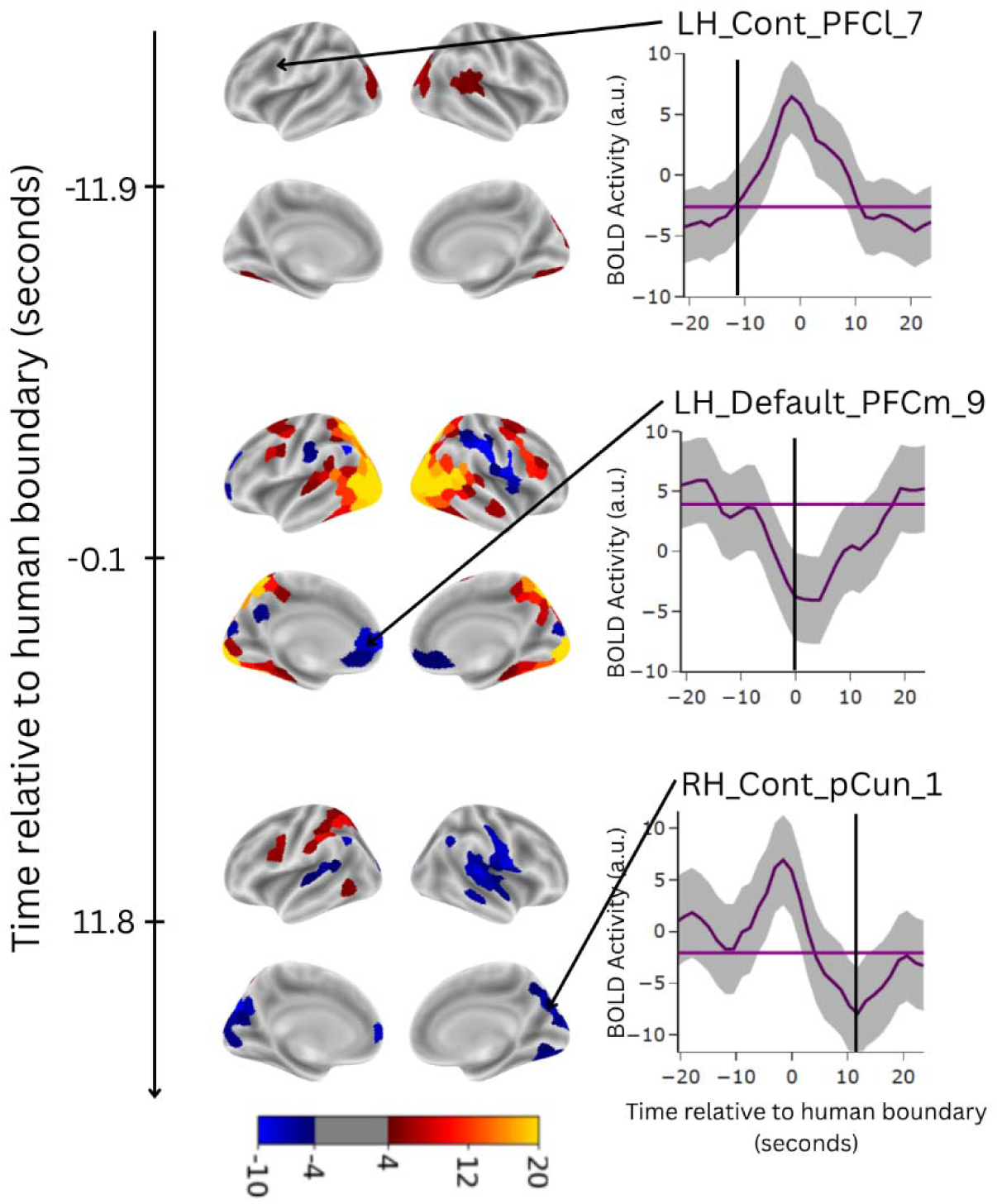
Brain activity relative to estimated baseline activity -11.9s before event boundary (top), at event boundary (middle), and 11.8s after event boundary (bottom). Brain maps display BOLD signal changes across multiple networks, with warm colors (red/yellow) indicating increased activity and cool colors (blue) indicating decreased activity. Visual, dorsal attention, and control network parcels tend to show increased BOLD activity around event boundaries, while default network regions exhibit decreased activity. Time courses (right panels) illustrate activity patterns in specific parcels (representative timecourses observed) within control network and default network (shades indicate plus and minus 2 standard errors). Parcel labels are from the Schaefer 400×7 parcellation atlas (Schaefer et al., 2018); for example LH_Cont_PFCl_7 indicates a parcel which is in the left lateral prefrontal cortex and is a component of the Control network. Some regions, notably the precuneus (bottom line graph), display a biphasic response characterized by initial activity increase at the boundary followed by subsequent decrease. Frame-by-frame brain maps and BOLD activity around boundaries for all parcels can be found at https://openneuro.org/datasets/ds005551 in the derivatives/figures/brain_maps_and_timecourses/ directory.

### Prefrontal and anterior temporal regions show early pattern shifts, followed by parietal shifts near boundaries, and whole-brain stabilization post-boundary

To model the effect of event boundaries on shifts in the local pattern of activity, we first calculated the amount of frame-to-frame change in the pattern of activity within each parcel and then fit FIR models to this pattern dissimilarity measure. Pattern dissimilarity was calculated as 1 - *r*, where *r* is the Pearson correlation between the voxel-wise activity patterns in the parcel for the two successive timepoints (see Methods for details). High pattern dissimilarity indicates that the neural patterns are more unstable (pattern shift), whereas low pattern dissimilarity indicates that neural patterns are more stable (pattern stabilization). As with the magnitude analyses, we fit the FIR model to a time window spanning approximately 20 seconds before and 20 seconds after human-identified boundaries for each parcel. This analysis revealed brain regions whose neural patterns shifted (increased dissimilarity) or stabilized (decreased dissimilarity) significantly (FDR with alpha < 0.001, corrected-p-value < 0.001) around event boundaries; these brain regions belong to the control (lateral PFC), default (medial PFC, posterior cingulate cortex, and temporal areas), dorsal attention (superior parietal lobule and posterior cortex) and visual (temporal occipital areas) networks (Figure 3).

**Figure 3:**
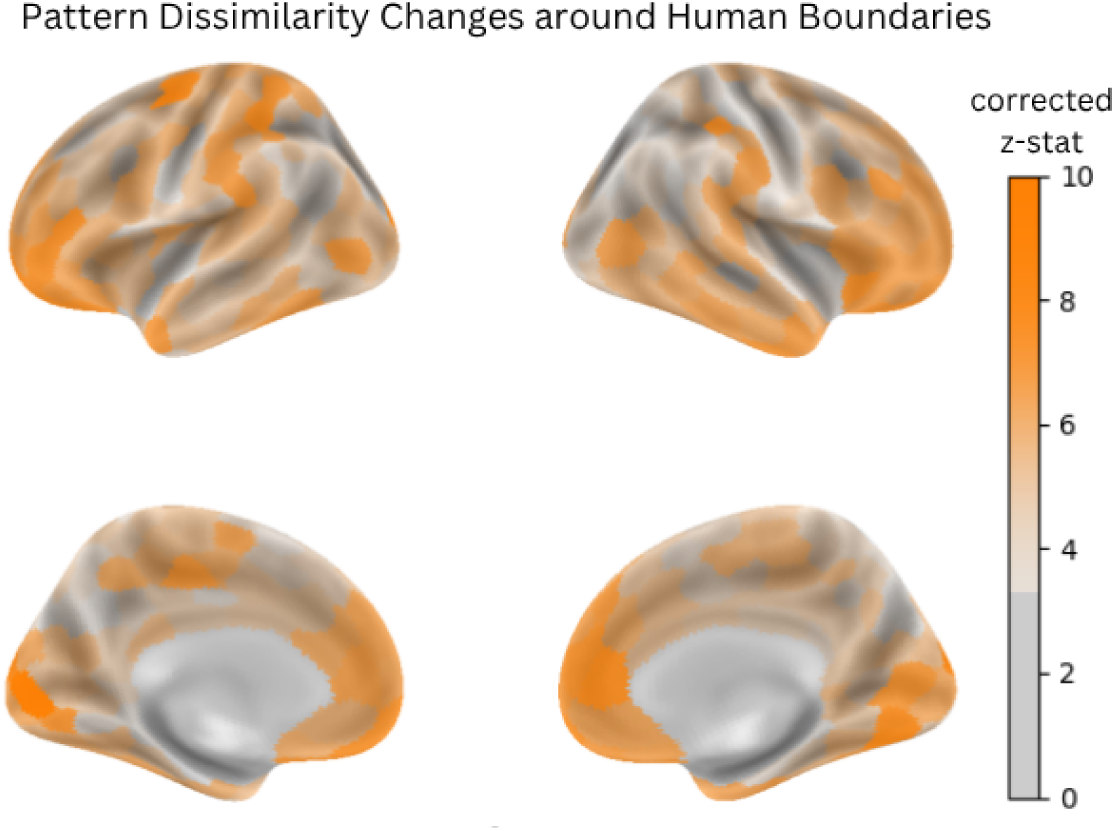
Regions where pattern dissimilarity changed significantly (corrected p < 0.001) around event boundaries. FIR models were used to predict pattern dissimilarity within a time window spanning approximately 20 seconds before and 20 seconds after human-identified event boundaries. Results show degree of changes in pattern dissimilarity relative to estimated baseline pattern dissimilarity in parcels across multiple brain networks, including the control network (lateral PFC), default network (medial PFC, posterior cingulate cortex, and temporal areas), dorsal attention network (superior parietal lobule and posterior cortex), and visual network (temporal occipital areas).

Next, we examined the same FIR model output for timecourses of pattern dissimilarity around event boundaries. Figure 4 shows changes in pattern dissimilarity relative to estimated baseline dissimilarity over time. The analysis revealed three salient phenomena. First, early pre-boundary temporal pole pattern shifts: At the early pre-boundary timepoint (-11.9s), pattern dissimilarity in the anterior temporal pole and orbitofrontal cortex significantly increased compared to estimated baseline dissimilarity, indicating pattern shifts in these regions. Second, immediate pre-boundary parietal pattern shifts: At the immediate pre-boundary timepoint (-4.5s), pattern dissimilarity in post central and superior parietal lobule areas (part of the dorsal attention network) increased significantly compared to estimated baseline dissimilarity, reflecting notable pattern shifts; moreover, the magnitude of immediate pre-boundary pattern shifts (-4.5s) is stronger than that of pattern shifts at early pre-boundary timepoint (-11.9s). Third, post-boundary whole-brain pattern stabilization: At the post-boundary timepoint (+11.8s), pattern dissimilarity across the whole brain decreased the most compared to estimated baseline dissimilarity, indicating pattern stabilization. This period of stability may represent the establishment and maintenance of a new event model. The result that patterns started to stabilize after boundaries and reached their peak of stabilization after about 10s suggest that the brain might take several seconds to build a new stable representation of the environment. The widespread nature of this stabilization suggests a global reconfiguration of brain states, consistent with theories proposing that event boundaries mark transitions between distinct neural states representing different events.

**Figure 4:**
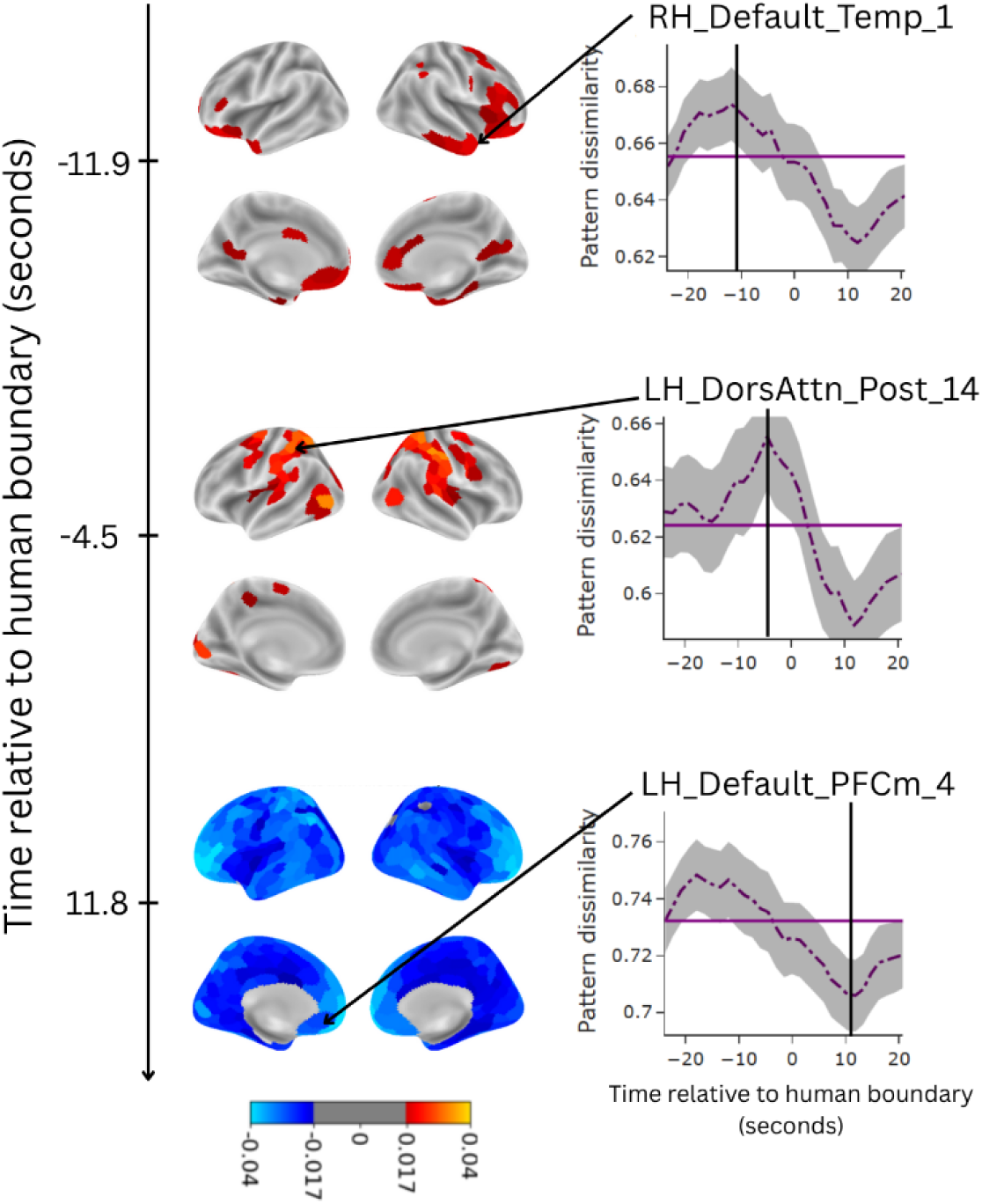
Event boundaries are characterized by increases in pattern dissimilarity, followed by post-boundary stability. Brain surface maps show deviations from baseline pattern dissimilarity (orange: increased dissimilarity/pattern shifts; blue: decreased dissimilarity/pattern stabilization) at three points in time. Line plots for selected regions show pattern dissimilarity for selected parcels over time, illustrating the representative timecourses observed (shades indicate plus and minus 2 standard errors). Baseline dissimilarity levels are indicated by horizontal purple lines. Higher values reflect greater pattern shifts, while lower values indicate more stable patterns. Parcel labels followed the Schaefer 400×7 parcellation atlas (Schaefer et al., 2018). Three key phenomena emerge: (1) At early pre-boundary (-11.9s), anterior temporal pole regions and orbitofrontal cortex showed significantly increased dissimilarity relative to baseline dissimilarity, indicating pattern shifts; (2) At immediate pre-boundary (-4.5s), superior parietal lobule and postcentral areas within the dorsal attention network exhibited increased pattern dissimilarity, indicating pattern shifts; (3) At post-boundary (+11.8s), pattern dissimilarity across the whole brain decreased, indicating widespread pattern stabilization. Brain maps and pattern dissimilarity around boundaries for all parcels can be found at https://openneuro.org/datasets/ds005551 in the derivatives/figures/brain_maps_and_timecourses/ directory.

The sequence of changes we observed—from temporal areas to parietal regions and finally to global stabilization—may reflect different stages of event model updating. The engagement of multiple brain regions at different time points also suggests that event segmentation may involve several distinct cognitive processes, each potentially implemented by different brain networks. This possibility sets the stage for our subsequent investigation of two potential event model updating control mechanisms: error-driven updating and uncertainty-driven updating. By examining the neural correlates of error-driven and uncertainty-driven updating separately, we can potentially identify dissociable neural mechanisms underlying human event segmentation.

### Error-driven boundaries and uncertainty-driven boundaries uniquely predict human boundaries

To investigate error-driven and uncertainty-driven event segmentation, we developed two computational models—one in which event model updating is triggered by an increase in prediction error and another in which event model updating is triggered by an increase in prediction uncertainty (Nguyen et al., 2024). The models generated error-driven or uncertainty-driven boundaries (see Methods). Discrete human and model event boundaries were transformed into boundary densities by applying a Gaussian density kernel (bandwidth = 4.45 seconds). Model-derived boundary densities were then employed to jointly predict human boundary density (see Methods), an analysis not performed in the previous work (Nguyen et al., 2024). This analysis revealed that boundary densities from both uncertainty-driven and error-driven models uniquely predicted human boundary density (t-statistics are 14.32 and 5.14 respectively, p < .001, df = 1636), even though error-driven boundary density is correlated with uncertainty-driven boundary density (mean correlation: 0.499, 95% CI: [0.401, 0.607]). This finding suggests that humans might rely on both types of prediction quality signals – errors in prediction and uncertainty about predictions – when segmenting events in continuous experience. Quantitatively, the combination of error-driven and uncertainty-driven boundaries explained approximately 20% of the variance in human-identified boundaries. To contextualize this result, human behavioral boundary identification typically has a reliability of approximately 0.6, which sets an upper limit of 36% for explainable variance. The relatively high explained variance suggests that our model-estimated error-driven and uncertainty-driven boundaries are good candidates for representing human error-driven and uncertainty-driven event updating processes. Model boundary densities and human boundary densities for four videos are depicted in Figure 5; boundary density peaks were used for all Finite Impulse Response analyses (see Methods). Based on these findings, we proceeded to use our model-estimated error-driven and uncertainty-driven boundaries as proxies for human cognitive processes in subsequent analyses. This approach allows us to investigate the neural correlates of error-driven and uncertainty-driven event segmentation separately, potentially revealing distinct brain networks associated with these two complementary processes.

**Figure 5:**
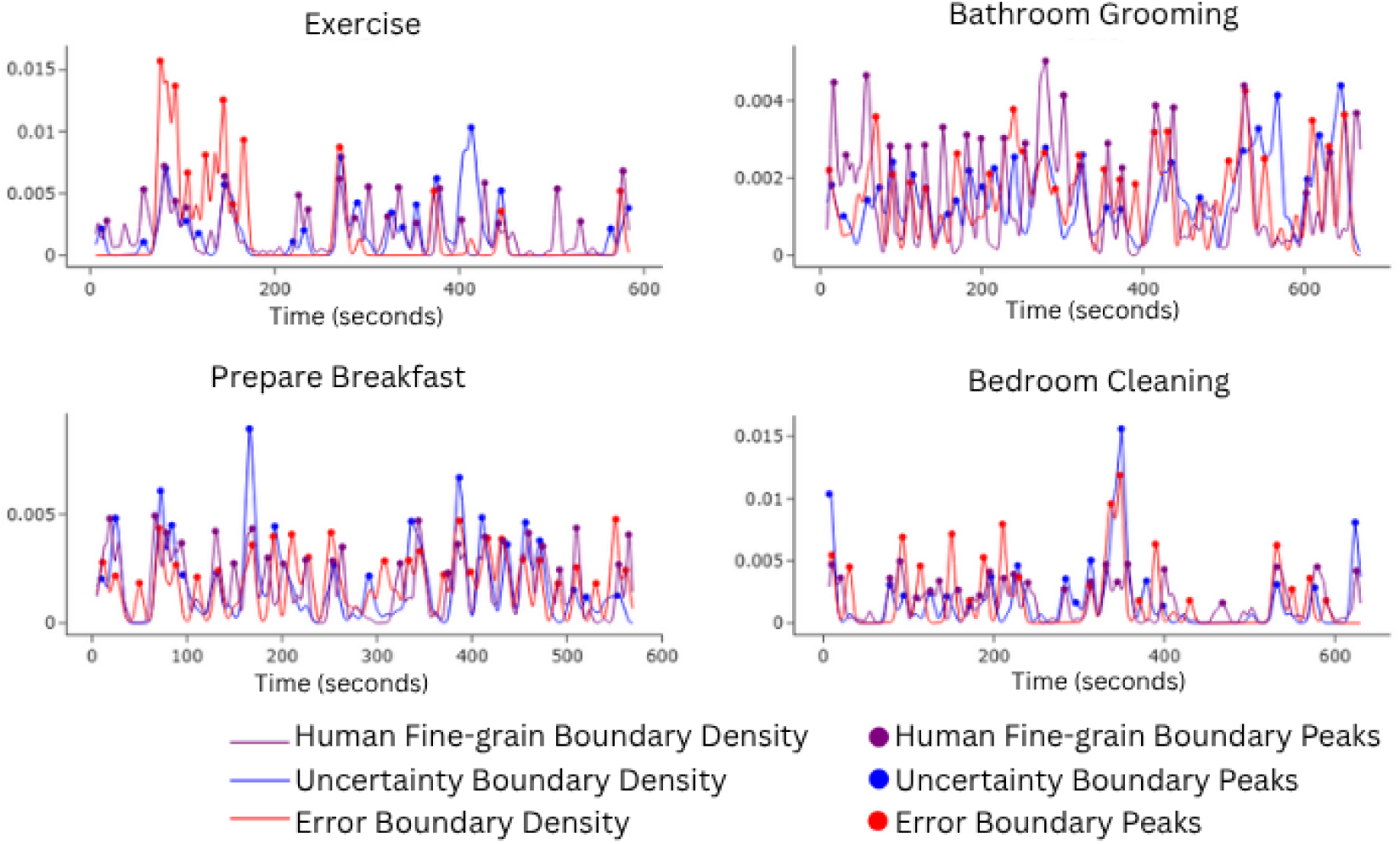
Error-model, uncertainty-model, and human boundary density and peaks in the four movie stimuli. Error-model boundary density and uncertainty-model boundary density both uniquely predict human boundary density. Boundary peaks were used for all FIR analyses.

### Error-driven and uncertainty-driven boundaries are associated with distinct brain networks and temporal dynamics

Building on the observation that human-identified boundaries are associated with pattern shifts in prefrontal, parietal, and temporal regions, and on our hypothesis that these regions might be responsible for distinct event model updating mechanisms, we sought to disentangle the neural correlates of error-driven and uncertainty-driven event model updating. Our first aim was to identify whether two qualitatively different types of boundaries are linked to pattern shifting and stabilization in overlapping or distinct brain areas. Thus, for each brain parcel, we fitted two FIR models to predict pattern dissimilarity: one using only error-driven boundaries (error FIR) and another using only uncertainty-driven boundaries (uncertainty FIR), and compared their performance.

This analysis revealed a dissociation between brain regions showing significant (FDR with alpha < 0.001, corrected-p-value < 0.001) neural pattern shift or stabilization around error-driven versus uncertainty-driven boundaries (Figure 6). Regions exhibiting strong neural pattern shift or stabilization around error-driven boundaries included the ventrolateral PFC (Figure 6, reddish regions). Regions exhibiting strong neural pattern shift or stabilization around uncertainty-driven boundaries included the postcentral gyrus, particularly those associated with the dorsal attention network, areas in the mid cingulate cortex associated with the ventral attention network, and areas within the visual network (Figure 6, blueish regions). Regions exhibiting strong neural pattern shift or stabilization around both types of boundaries included the medial PFC and temporal components of the default network (Figure 6, magenta regions). These results reveal distinct brain networks whose neural pattern shifted or stabilized around error-driven or uncertainty-driven boundaries, each partially aligning with regions showing neural pattern shift or stabilization around human boundaries (Figure 3). The identification of two dissociable sets of brain regions associated with each boundary type, and the finding that both sets partially overlapped with regions involved in human-identified boundaries, suggests that both error-driven and uncertainty-driven signals play roles in event model updating. This neural evidence corroborates and extends the behavioral findings that both types of boundaries uniquely predict human-identified boundaries. To account for the correlation between uncertainty-driven boundaries and error-driven boundaries, we also fitted a FIR model that predicted pattern dissimilarity from both types of boundaries (combined FIR) for each parcel. Then, we performed two likelihood ratio tests: combined FIR to error FIR, which measures the unique contribution of uncertainty boundaries to pattern dissimilarity, and combined FIR to uncertainty FIR, which measures the unique contribution of error boundaries to pattern dissimilarity. The analysis also revealed two dissociable sets of brain regions associated with each boundary type (see Figure S1).

**Figure 6:**
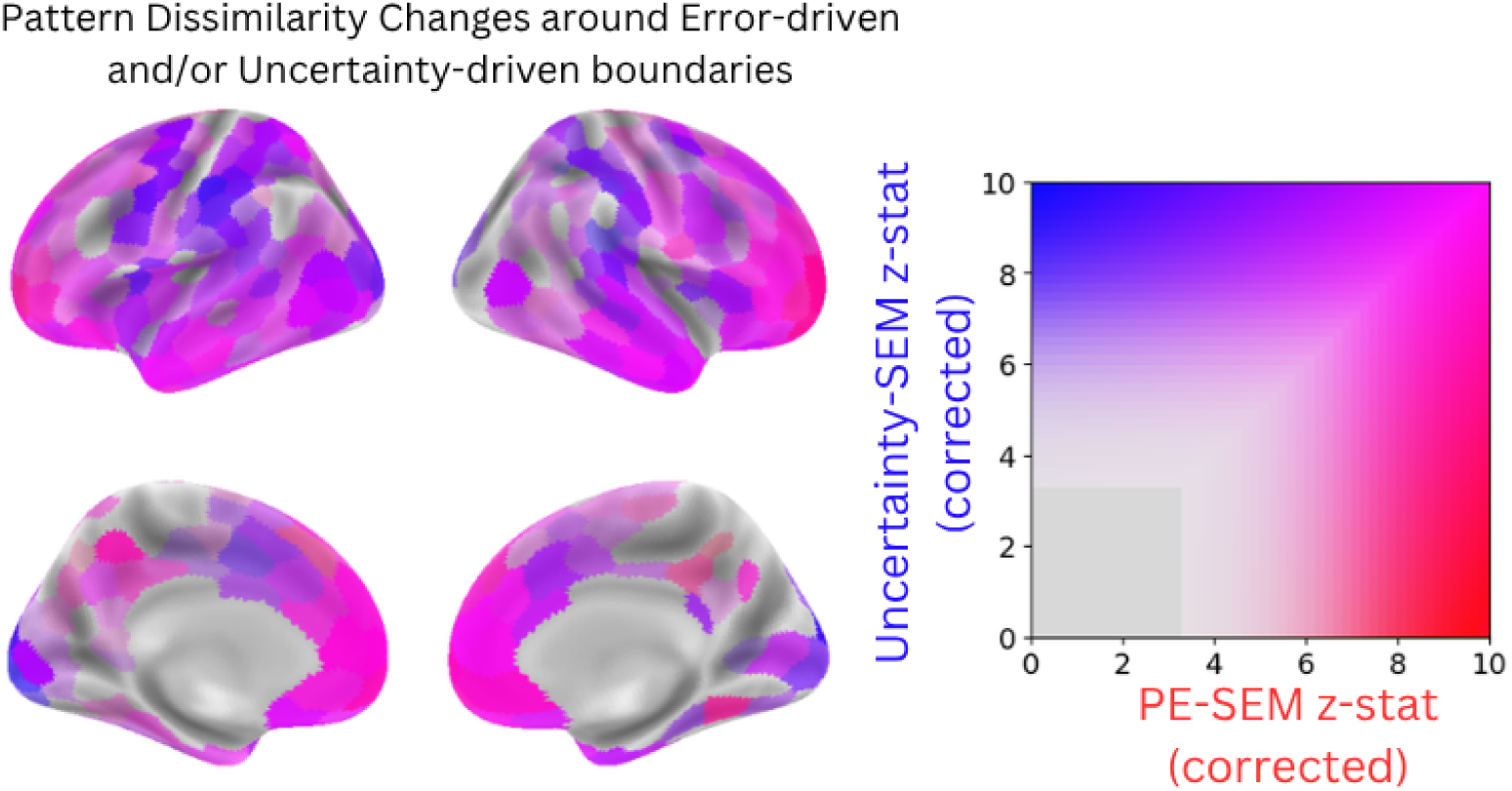
Regions showing significant (corrected p < 0.001) changes in pattern dissimilarity relative to estimated baseline dissimilarity for error-driven and/or uncertainty-driven boundaries. Regions in red show stronger changes in pattern dissimilarity relative to estimated baseline pattern dissimilarity for error-driven boundaries, while regions in blue show stronger changes in pattern dissimilarity relative to estimated baseline pattern dissimilarity for uncertainty-driven boundaries, and regions in magenta show strong changes in pattern dissimilarity for both boundary types.

Next, we used the two FIR models to investigate the time-course of changes in pattern dissimilarity around error-driven and uncertainty-driven boundaries. This analysis is crucial for understanding the temporal sequence of cognitive processes involved in event segmentation and how they might differ between these two mechanisms. It revealed distinct spatial and temporal changes in pattern dissimilarity, which partially overlap with the changes observed for human-identified boundaries discussed previously. Some regions showed early pre-boundary (-11.9s) neural pattern shifts: For error-driven boundaries, increased pattern dissimilarity was observed in the ventrolateral PFC, which is part of the control network (Figure 7A, top). For uncertainty-driven boundaries, we observed increased pattern dissimilarity (indicating pattern shifts) in temporal areas and the PFC (Figure 7B, top). These early pattern shifts associated with error-driven and uncertainty-driven boundaries corresponded to subcomponents of the early shifts observed before human-identified boundaries (Figure 4, top). Another group of regions showed immediate pre-boundary (-4.5s) pattern shifts: error-driven boundaries were associated with increased pattern dissimilarity primarily in the left ventrolateral PFC and anterior temporal pole (Figure 7A, middle). Uncertainty-driven boundaries were associated with increased pattern dissimilarity in parietal areas, occipital areas, temporal areas, and prefrontal areas, with strong shifts in regions within the dorsal attention network (Figure 7B, middle).

**Figure 7:**
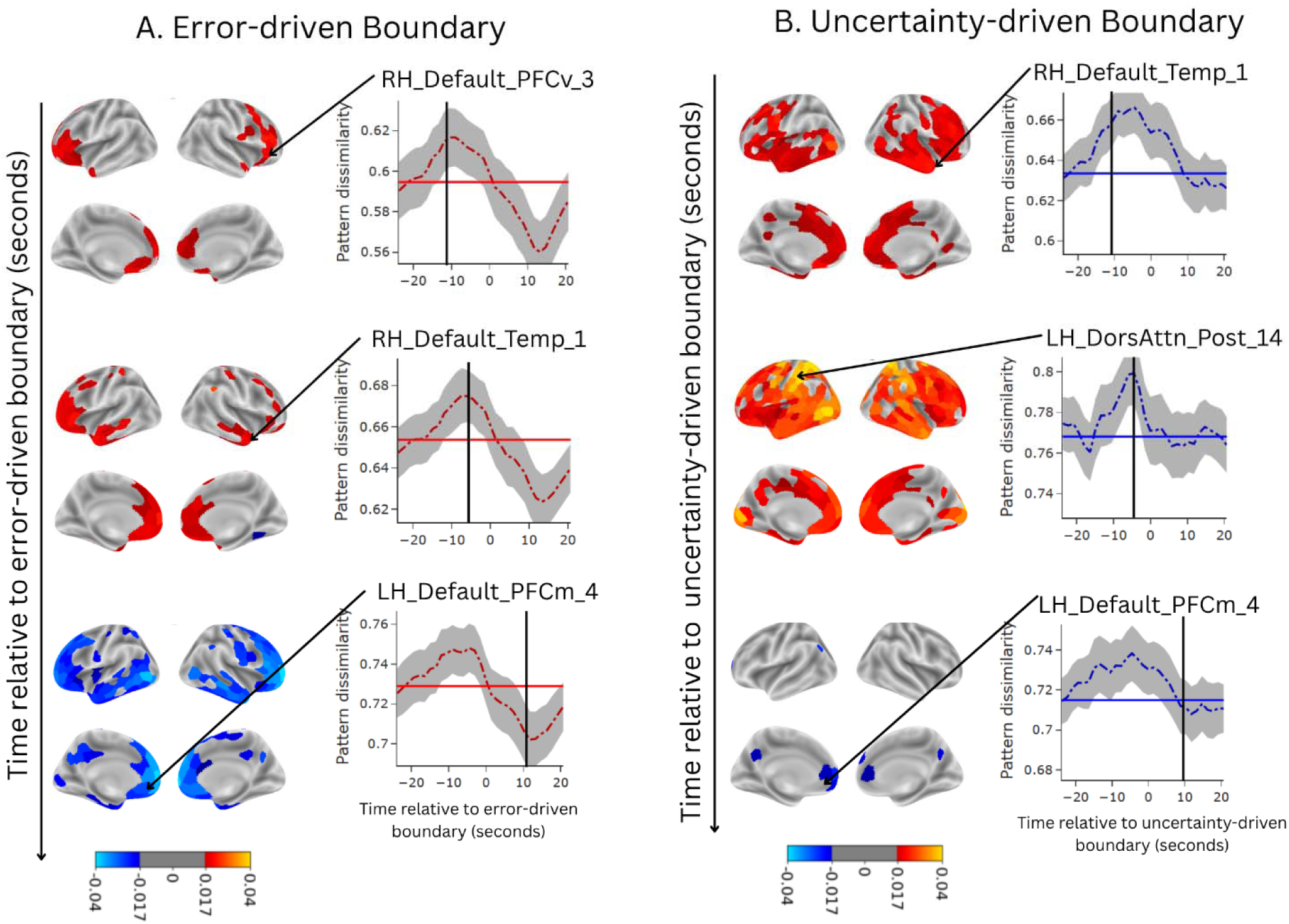
Changes in pattern dissimilarity relative to baseline dissimilarity around boundaries identified by (A) error-driven model and (B) uncertainty-driven model. Brain surface maps show deviations from baseline pattern dissimilarity (orange: increased dissimilarity/pattern shifts; blue: decreased dissimilarity/pattern stabilization). Line plots for selected regions show pattern dissimilarity over time, with baseline dissimilarity levels indicated by horizontal blue (uncertainty-driven) or red (error-driven) lines (shades indicate plus and minus 2 standard errors). Higher dissimilarity values reflect greater pattern shifts, while lower dissimilarity values indicate more stable patterns. Parcel labels followed the Schaefer 400×7 parcellation atlas (Schaefer et al., 2018). At early pre-boundary (-11.9s), error-driven boundaries corresponded to increased pattern dissimilarity primarily in ventrolateral PFC, while uncertainty-driven boundaries corresponded to more widespread pattern shifts in temporal areas, dorsomedial and dorsolateral prefrontal regions, and anterior temporal cortex. At immediate pre-boundary (-4.5s), error-driven boundaries corresponded to pattern shifts predominantly in the left ventrolateral PFC and anterior temporal pole, whereas uncertainty-driven boundaries corresponded to more extensive pattern shifts across parietal, occipital, temporal, and prefrontal areas, particularly strong within the dorsal attention network. At post-boundary timepoints (+11.8s), error-driven boundaries corresponded to widespread pattern stabilization (decreased dissimilarity) across prefrontal, temporal, and occipital areas with strongest effects in prefrontal cortex, while uncertainty-driven boundaries show limited pattern stabilization restricted primarily to portions of the medial PFC. Brain maps and pattern dissimilarity around error-driven boundaries or uncertainty-driven boundaries for all parcels can be found at https://openneuro.org/datasets/ds005551 in the derivatives/figures/brain_maps_and_timecourses/ directory. Brain maps and neural activity around error-driven and uncertainty-driven boundaries for all parcels can also be found in the same directory.

These pattern shifts for uncertainty-driven boundaries corresponded to our second observation from human boundaries, for which parietal regions showed significant pattern shifts (Figure 4, middle). Moreover, pattern shifts for both error-driven boundaries and uncertainty-driven boundaries were higher and involve more brain areas at immediate pre-boundary (-4.5s) compared to early pre-boundary (-11.9s). Finally, many regions showed post-boundary (+11.8s) pattern stabilization: error-driven boundaries were followed by widespread decreases in pattern dissimilarity in prefrontal, temporal, and occipital parcels, with strongest stabilization in the prefrontal cortex (Figure 7A, bottom). The pattern stabilization following error-driven boundaries corresponds with our third observation from human boundaries, where pattern stabilization was strongest for the PFC (Figure 4, bottom). For uncertainty-driven boundaries, we only observed decreased pattern dissimilarity in part of the medial PFC (Figure 7B, bottom).

The above analysis revealed shared brain regions whose changes in pattern dissimilarity were associated with both error-driven and uncertainty-driven boundaries (magenta regions in Figure 6). Although these two boundary types evoked neural pattern shift or stabilization in overlapping brain regions, they may drive distinct temporal dynamics of pattern shift and stabilization. To test this hypothesis, for each parcel that showed significant changes in pattern dissimilarity for both boundary types (FDR with alpha < 0.001, corrected-p-value < 0.001 for both error-driven and uncertainty-driven FIR models), we computed correlations between pattern dissimilarity timecourses for error-driven and uncertainty-driven boundaries and compared them against a null distribution. This null distribution was constructed from within-boundary-type correlations, using bootstrapped timecourses for both error-driven and uncertainty-driven boundaries separately. The resulting z-statistic maps (Figure 8) revealed statistically significant (FDR with alpha < 0.001, corrected-p-value < 0.001) differences in temporal dynamics between the two boundary types, particularly in prefrontal and temporal regions. For example, two parcels in the left medial PFC (LH_Default_PFCm_4) and in the right anterior temporal pole (RH_Default_Temp_1), whose changes in pattern dissimilarity were associated with both error-driven and uncertainty-driven boundaries, had different temporal dynamics for each boundary type (Figure 7A middle and 7B top). This result suggests that even though error-driven and uncertainty-driven event model updating were associated with the same brain regions, the two updating mechanisms were associated with distinct temporal dynamics of pattern shift and stabilization in these shared regions.

**Figure 8:**
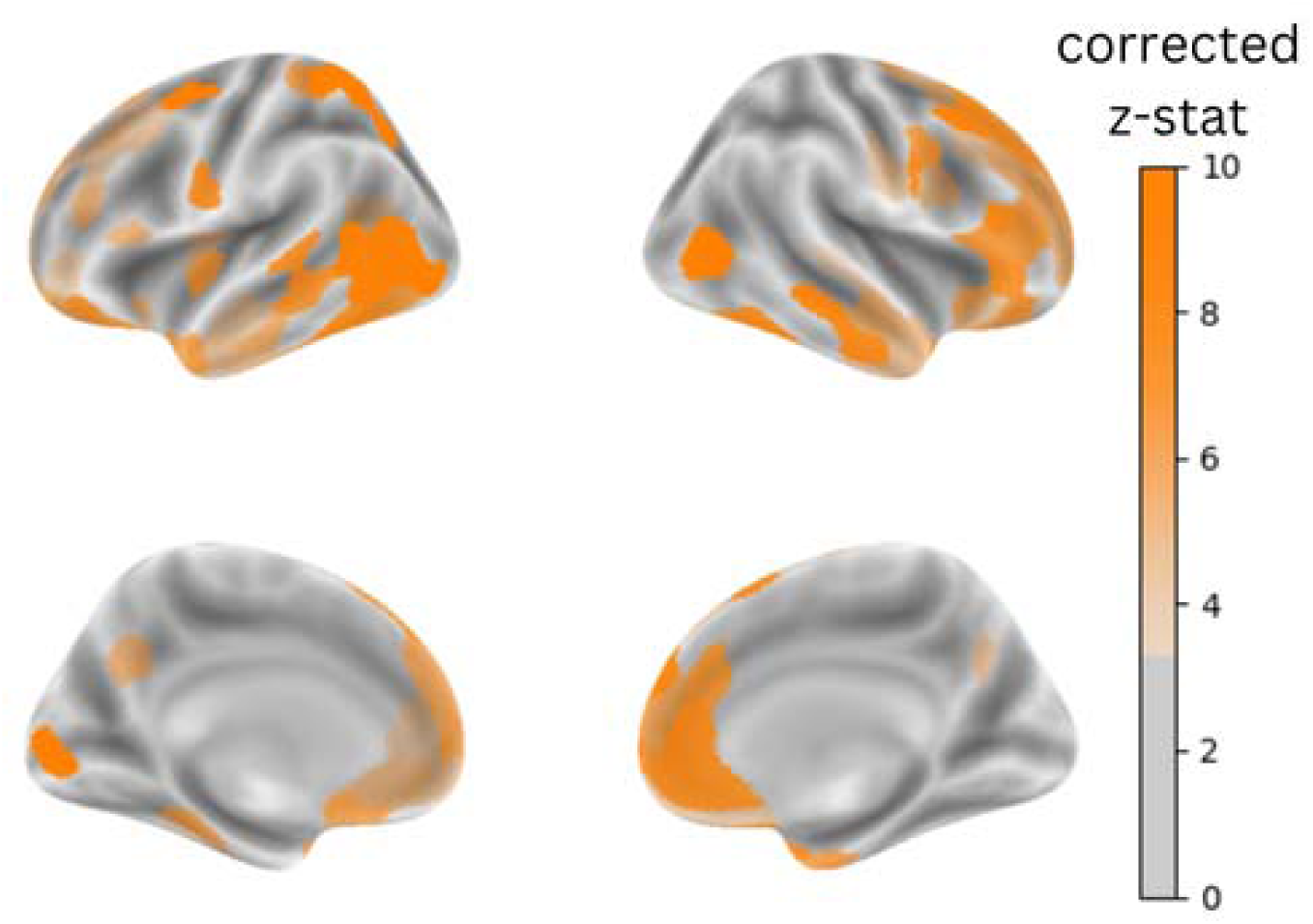
Comparison of pattern dissimilarity timecourses between error-driven and uncertainty-driven boundaries. Z-statistics maps show regions where pattern dissimilarity timecourses differ between the two boundary types, with higher values (more orange) indicating stronger differences in temporal dynamics. The most pronounced differences observed in the prefrontal cortex, posterior parietal regions, and portions of the temporal cortex.

## Discussion

Humans and other animals navigate a world that is complex and dynamic, but that is also characterized by stable dynamical regimes during which sequential activity is predictable. To cope with the complexity of experience and to leverage the intervals of predictability, the nervous system constructs stable models of the current situation on a hierarchy of timescales from seconds to tens of minutes (Baldassano et al., 2017; Hasson et al., 2015; Richmond & Zacks, 2017). The present study addressed a central question about these event models: What control mechanism determines when they are updated?

We first characterized the neural signatures of human event segmentation by examining both univariate activity changes and multivariate pattern changes around subjectively identified event boundaries. Using multivariate pattern dissimilarity, we observed a structured progression of neural reconfiguration surrounding human-identified event boundaries. The largest pattern shifts were observed near event boundaries (∼4.5s before) in dorsal attention and parietal regions; these correspond with regions identified by Geerligs et al. as shifting their patterns on a fast to intermediate timescale (2022). We also observed smaller pattern shifts roughly 12 seconds prior to event boundaries in higher-order regions within anterior temporal cortex and prefrontal cortex, and these are slow-changing regions identified by Geerligs et al. (2022). This is puzzling. One prevalent proposal, based on the idea of a cortical hierarchy of increasing temporal receptive windows (TRWs), suggests that higher-order regions should update representations after lower-order regions do (Chang et al., 2021). In this view, areas with shorter TRWs (e.g., word-level processors) pass information upward, where it is integrated into progressively larger narrative units (phrases, sentences, events). This proposal predicts neural shifts in higher-order regions to follow those in lower-order regions. By contrast, our findings indicate the opposite sequence. Our findings suggest that the brain might engage in top-down event representation updating, with changes in coarser-grain representations propagating downward to influence finer-grain representations. (Friston, 2005; Kuperberg, 2021). For example, in a narrative where the main goal is achieved midway—such as a detective solving a mystery before the story formally ends—higher-order regions might update the overarching event representation at that point, and this updated model could then cascade down to reconfigure how lower-level regions process the remaining sensory and contextual details. In the period after a boundary (around +12 seconds), we found widespread stabilization of neural patterns across the brain, suggesting the establishment of a new event model. Future work could focus on understanding the mechanisms behind the temporal progression of neural pattern changes around event boundaries.

We then took a computational cognitive neuroscience approach to disentangle the contributions of two potential updating mechanisms—prediction error and prediction uncertainty—in driving event model updating. Building on the theoretical framework proposed by Baldwin and Kosie (2021), we developed two computational models to generate boundaries driven by prediction error or prediction uncertainty. Both models uniquely explained variance in human boundary judgments. Together, these model-derived boundaries explained more than half of the explainable variance in human segmentation. This result provides evidence that both prediction error and prediction uncertainty contribute to how observers parse ongoing experience into events. More importantly, error-driven and uncertainty-driven boundaries were associated with distinct brain networks exhibiting different temporal dynamics. Error-driven boundaries were linked to early neural pattern shifts in ventrolateral prefrontal areas, followed by pronounced pattern stabilization in prefrontal and temporal regions. In contrast, uncertainty-driven boundaries were associated with pattern shifts in across wide areas of the brain, especially in parietal regions within the dorsal attention network, and with minimal subsequent stabilization in part of the medial PFC. Interestingly, both boundary types evoked pattern shifts in overlapping regions within the default network, but with different temporal dynamics. The dissociations in both spatial and temporal neural signatures provide evidence for the theoretical distinction proposed by Baldwin and Kosie (2021), that humans rely on both prediction error and prediction uncertainty to parse continuous activities into meaningful events. Our findings suggest that event segmentation is supported by two separate but complementary brain networks, each responsive to a different prediction quality signal. This dual-network architecture represents a significant advancement in our understanding of event segmentation, extending the claim of EST that declines in prediction quality trigger event model updating (Zacks et al. 2007) to encompass these two aspects of prediction quality: accuracy and certainty.

Previous studies using HMMs (Baldassano et al., 2017) and GSBS (Geerligs et al., 2021, 2022) have identified neural state boundaries in multimodal regions that align closely with subjectively perceived event boundaries. Consistent with these findings, our multivariate analyses also revealed pattern shifts around human boundaries in regions previously identified, including areas within the default mode network (medial prefrontal cortex, posterior cingulate cortex, and anterior temporal regions) and dorsal attention network. Despite the advances made in this study, several limitations and open questions remain. Although our computational models successfully captured a substantial portion of variance in human boundary judgments, approximately 16% of the explainable variance remains unaccounted for. This suggests that additional mechanisms beyond prediction error and prediction uncertainty may contribute to event segmentation. One candidate mechanism comes from latent cause inference models (Kuperberg, 2021; Shin & DuBrow, 2021). Whereas Zacks et al. (2007) and Baldwin and Kosie (2021) assume the cognitive system maintains an event model and uses prediction error or prediction uncertainty from that model to segment events, latent cause inference models start from a different assumption. These models posit that the cognitive system monitors a probability distribution over unobservable latent causes that are most likely to generate current sensory observations. Event boundaries in this framework correspond to moments of high uncertainty over which latent cause is currently active. Future studies could develop computational models based on latent cause inference principles and compare their neural signatures with those of error-driven and uncertainty-driven boundaries identified in our study.

Although our error-driven and uncertainty-driven models segmented and categorized continuous experience in a manner that aligns with human behavior (Nguyen et al., 2024), the correspondence is far from perfect. Consequently, the event boundaries identified by these two models might not fully capture the actual error-driven and uncertainty-driven processes that humans employ. The current evidence for more than one model updating mechanism opens the door for an architecture in which a handful of different triggers may govern different components of event model updating. Such an architecture would be inelegant—but evolution’s engineering solutions are sometimes complex and kludgy rather than simple and sleek.

In conclusion, this study provides behavioral and neural evidence for two overlapping brain networks that maintain and update representations of the environment, controlled by either prediction error or prediction uncertainty. Error-driven boundaries were uniquely associated with representational shifts in ventrolateral prefrontal cortex and subsequent pattern stabilization in temporal and prefrontal areas. Uncertainty-driven boundaries were uniquely linked to representational shifts in dorsal attention network with minimal subsequent stabilization. Both boundary types were associated with pattern shifts in regions within the default mode network, such as medial PFC and temporal areas, which have previously been hypothesized to maintain and update event representations. By identifying the neural correlates of theoretically distinct updating mechanisms, these results form a basis for an integrative account of the computational and neurophysiological aspects of how the brain maintains stable representations of complex everyday activity.

## Methods

### Human and model segmentation

Human event boundaries for four activities were obtained from a norming study in which 30 online participants (from Mechanical Turk) watched each activity and marked event boundaries (Bezdek et al., 2022). Error-driven boundaries and uncertainty-driven boundaries were generated by two computational models developed previously (Nguyen et al., 2024). Both models maintain an active event representation that continuously predicts how activity will unfold. The error-driven model triggers event boundary detection when prediction error— operationalized as the Euclidean distance between observed and predicted scene vectors— exceeds a threshold. In contrast, the uncertainty-driven model triggers event boundary detection based on prediction uncertainty, conceptualized as epistemic uncertainty that reflects the model’s confidence in its own predictions given its current knowledge state. This prediction uncertainty is computed by generating 32 different predictions using random dropout on the model weights and measuring the variance across these predictions (Gal & Ghahramani, 2016; Kendall & Gal, 2017). When their respective thresholds are exceeded, each model initiates an inference process to evaluate alternative event representations. Event boundaries are identified at moments when this inference process results in a switch from one event representation to another, reflecting the model’s decision that the current event representation no longer adequately captures ongoing activity. To account for model random initialization, we ran 16 simulations for each model, and we used event boundaries from all 16 simulations. The error-driven and uncertainty-driven models were tuned such that their median number of boundaries were as close as possible to human fine-grain median number of boundaries (Nguyen et al., 2024). To analyze the relationships between the three kinds of boundaries, we applied a Gaussian kernel density function (bandwidth = 4.45 seconds) on the human, error-driven, and uncertainty-driven event boundaries to obtain continuous event boundary densities. In later analyses, error-driven and uncertainty-driven boundary densities were used as continuous predictors of the normed human boundary density.

To model the effects of human and model event boundaries on fMRI magnitude and pattern dissimilarity, we identified normative boundaries by selecting peaks in the event boundary densities. To identify peaks from continuous boundary densities, we ran a greedy algorithm to iteratively select one peak at a time with the highest density value, with a condition that the current peak is more than 7 seconds away from already selected peaks. The number of peaks for each movie was set to the median number of boundaries identified by the Bezdek (2022) study participants for each movie (Figure 5). These peaks were used in the finite impulse response analyses as reference points, such that each “trial” in the analysis represented a window around the event boundary (see below).

### Imaging Session

#### fMRI Participants

Imaging data were collected from 47 participants (age mean=23.3 years, SD=4.3, range 18-35; 32 female, 15 male). Their self-reported demographic groupings were 14 Asian, 2 Black/African-American, 26 white (5 Hispanic/Latino), and 5 More Than One Race. Imaging data from two additional participants (both white males, one Hispanic/Latino, 19 and 23 years) could not be obtained (equipment failure, incidental finding) and are not included here. All participants were right-handed, fluent English speakers, and without MRI safety contraindications or neurological impairments. Participants provided written informed consent in accordance with the Institutional Review Board at Washington University in St. Louis, and received $125 compensation for completion of all sessions.

#### Acquisition parameters

All scans were acquired using a 3T Siemens Prisma with a 64-channel head coil in the East Building MR Facility of the Washington University Medical Center, between May 2021 and May 2022. Imaging files were sent from the scanner to the Washington University Central Neuroimaging Data Archive (Gurney, 2017) XNAT (RRID:SCR_003048) database for permanent archiving. FIRMM (Dosenbach et al., 2017) was used to monitor participant motion throughout the imaging session. Scan duration was not modified by FIRMM, but if it indicated excessive movement the experimenter notified the participant and repositioned them, as needed.

Each imaging session began with T1 and T2-weighted high-resolution ABCD MPRAGE+PMC structural scans (T1: 1 mm isotropic voxels; 2.5 s TR, 2.9 msec TE, 1.07 s TI, 8° flip angle; T2: 1 mm isotropic voxels; 3.2 s TR, 0.565 s TE, 120° flip angle), followed by four functional task runs. The functional (BOLD, blood oxygenation level dependent) scans were acquired with CMRR multiband sequences (University of Minnesota Center for Magnetic Resonance Research) (Feinberg et al., 2010; Xu et al., 2013), multiband (simultaneous multislice; SMS) factor 4, without in-plane acceleration (iPat = none), resulting in 2.08 mm isotropic voxels, 1.483 s TR (TE 37 msec, flip angle 52°; protocol sheets at https://openneuro.org/datasets/ds005551 under derivatives/ Scanning_parameters_NP1157new_May8_2021.pdf). All functional runs were collected with Anterior to Posterior (“AP”) encoding direction, and proceeded by a pair (AP, PA) of spin echo field maps and a single-band reference (“SBRef”) image.

#### Task runs

All participants completed four task fMRI runs, each of which consisted of watching a silent video. The four video stimuli (and accompanying normative event boundaries used for later analyses) are from the Multi-angle Extended Three-dimensional Activity (META) stimulus set (Bezdek et al., 2022). Each video depicts a different actor carrying out different everyday activities (exercising, cleaning a room, making breakfast, bathroom grooming), all filmed from a fixed camera position.

A different video was shown in each fMRI run, with all participants viewing the same four videos in the same order (Table 1). Participants were instructed, “As you watch, please pay attention and try to remember what occurs, as your memory will be tested later.” A brief (8-11 second) fixation cross proceeded the start of each film and reappeared when the film completed.

**Table 1:** Video stimuli characteristics and fMRI scanning parameters for each run.

|  | Run 1 | Run 2 | Run 3 | Run 4 |
| --- | --- | --- | --- | --- |
| Video file name | 1.2.3_C1_trim.mp4 | 6.3.9_C1_trim.mp4 | 3.1.3_C1_trim.mp4 | 2.4.1_C1_trim.mp4 |
| Video file location | https://osf.io/7hqr9 | https://osf.io/73yqe | https://osf.io/pdg43 | https://osf.io/bm589 |
| Video duration | 9:44 min | 11:14 min | 9:43 min | 10:34 min |
| fMRI run duration | 405 TRs; 10 min | 467 TRs; 12 min | 404 TRs; 10 min | 444 TRs; 11 min |

#### Image processing

Image preprocessing was performed by *fMRIPrep* 20.2.3 (Esteban, Markiewicz, et al. (2018); Esteban, Blair, et al. (2018); RRID:SCR_016216), which is based on *Nipype* 1.6.1 (Gorgolewski et al. (2011); Gorgolewski et al. (2018); RRID:SCR_002502); boilerplate text at Supplemental Information. Both the preprocessed images (spatially normalized to the MNI152NLin2009cAsym template) and realignment parameters (_desc-confounds_timeseries.tsv) were used in later analyses. Image quality was evaluated by reviewing temporal mean and standard deviation images (Etzel, 2023) and deemed acceptable in all participants. Frames with more than 0.5 mm FD were censored. Participants were retained only if more than half their data was usable; two participants had two or more runs at the 20% censoring threshold and so were dropped from analysis. Quality control summary files are available at https://openneuro.org/datasets/ds005551 under derivatives/QCKnitrs/ directory.

Preprocessed fMRI data was further processed by regressing drifts (linear, polynomial, and cubic) and 6 motion confounds from every voxel BOLD timeseries. Next, a standard parcellation was used: the Schaefer 400 parcels (17-network labels) for cortex (Schaefer et al., 2018). All analyses were performed at the parcel-level. All voxel BOLD timeseries were shifted by 4 TR (approximately 5.93s) to account for hemodynamic lag.

### Parcel BOLD activity and parcel pattern dissimilarity

To quantify parcel BOLD activity we averaged across all voxels in every parcel, resulting in a single value for each parcel in each timepoint (TR) of the four functional runs.

To quantify parcel neural pattern shifts, we calculated temporal pattern dissimilarity for each brain parcel using the Schaefer 400-parcel atlas (Schaefer et al., 2018). For each timepoint (TR t), we computed the dissimilarity between patterns before and after that timepoint. Specifically, we first created stable pattern representations by averaging voxel activities across three consecutive TRs preceding the timepoint (TRs t-3, t-2, and t-1) and three consecutive TRs following the timepoint (TRs t+1, t+2, and t+3). This average approach reduces noise inherent in single-TR measurements. We then calculated the Pearson correlation between these two averaged patterns and subtracted the correlation value from 1 to obtain a dissimilarity measure.

### Finite impulse response analysis

For each parcel, we employed finite impulse response (FIR) analysis to examine the temporal dynamics of BOLD activity (average across voxels) and pattern shifts around event boundaries. The FIR model is a flexible extension of the general linear model (GLM) commonly used in fMRI analysis. Whereas GLM approaches to fMRI modeling often assume a fixed hemodynamic response function (HRF), the FIR model makes no assumptions about the shape of the response, instead estimating separate coefficients for each time point (lag) relative to event boundaries. The coefficients from the FIR model indicate changes relative to baseline, which can be conceptualized as the expected value when far from event boundaries. This approach is particularly valuable for investigating neural responses to event boundaries, where the temporal profile may differ from canonical HRF shapes and vary across brain regions. In our study, there were three types of boundary: human-annotated boundaries, and computationally derived boundaries(error-driven and uncertainty-driven models). We constructed separate FIR models for each boundary type, allowing us to compare how different types of event segmentation relate to brain responses in different parcels. This approach enabled us to distinguish between neural responses associated with computationally derived error-driven versus uncertainty-driven boundaries.

Individual FIR models were fit for each subject (first-level analysis) to estimate coefficients for each time lag relative to each type of boundary onsets. There were 10 time lags before the event boundary and 20 time lags after the event boundary, and successive time lags are 1.483 seconds apart (1 TR). FIR models were applied separately to both BOLD activity and pattern dissimilarity measures. After first-level analysis, we performed second-level analysis across subjects. Z-statistics were calculated by comparing the full FIR model to an intercept-only model that assumed no changes in BOLD activity or pattern dissimilarity in response to boundaries. These z-statistics were used to identify brain regions that showed significant BOLD activity changes and/or pattern shifts in response to different boundary types. Instead of considering only across-subject variance when estimating standard errors for coefficients, we incorporated both within-subject and across-subject variance components to better estimate the standard errors associated with coefficients at different lags. The custom R code to estimate both within-subject and across-subject variance is available at: https://openneuro.org/datasets/ds005551 under derivatives/scripts/fir_analyses.Rmd.

## Acknowledgements

The authors would like to thank Sarah Morse and Grace Zhou for fMRI data collection, Todd S. Braver for helpful feedback on data analyses, Hayoung Song for feedback on the manuscript, and the members of the Dynamic Cognition Laboratory for their thoughtful input.

## Additional information

## Funding

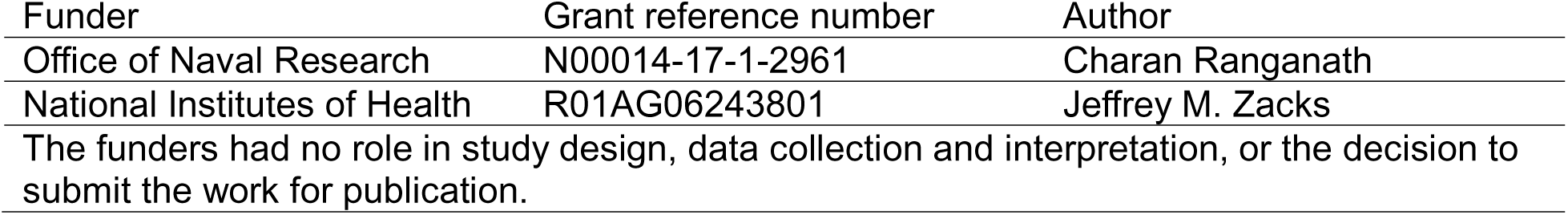

## Author Contributions

T.T.N.: conceptualization, data curation, formal analysis, investigation, methodology, visualization, writing—original draft, writing—review & editing. M.A.B.: conceptualization, data curation, formal analysis, investigation, methodology, visualization. J.E.B.: conceptualization, data curation, formal analysis, investigation, methodology, visualization, writing—review & editing. J.M.Z.: conceptualization, funding acquisition, project administration, supervision, writing—review & editing.

## Additional Files

### Data Availability

BIDS data for this project can be found at https://openneuro.org/datasets/ds005551 under sub-<PARTICIPANT_ID> directories.

All the data and analysis scripts used for this project can be found at https://openneuro.org/datasets/ds005551 under derivatives/scripts/ directory.

## Supplemental Information

### Error-driven and uncertainty-driven boundaries are associated with distinct brain networks and temporal dynamics

**Figure S1:**
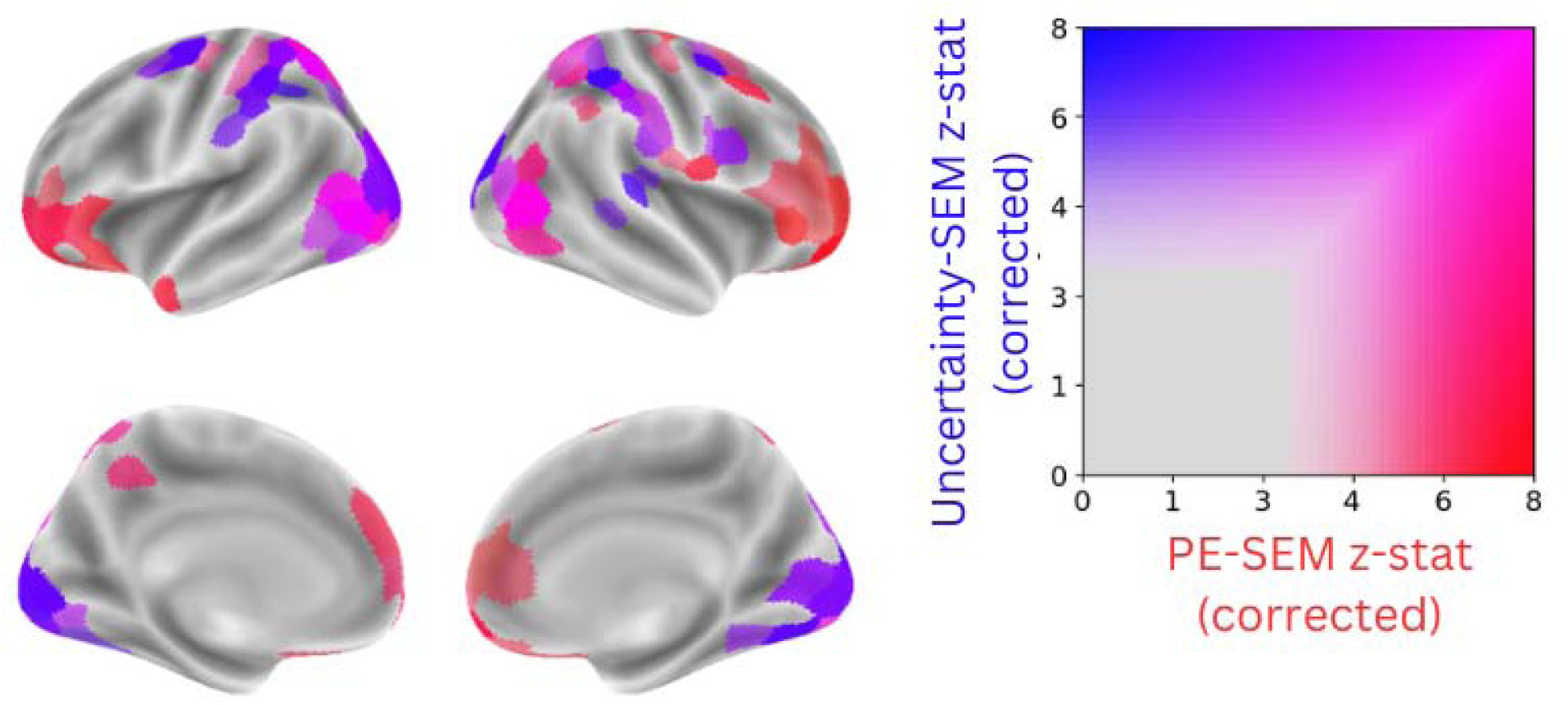
Regions showing unique contributions of error-driven and uncertainty-driven boundaries to changes in pattern dissimilarity (corrected p < 0.001). Unique effects were estimated using likelihood ratio tests comparing the combined FIR model with models including only one boundary type. Regions in red show stronger unique contributions of error-driven boundaries, while regions in blue show stronger unique contributions of uncertainty-driven boundaries, and regions in magenta show strong unique contributions of both boundary types.

#### fMRIprep boilerplate

##### Anatomical data preprocessing

A total of 1 T1-weighted (T1w) images were found within the input BIDS dataset. The T1-weighted (T1w) image was corrected for intensity non-uniformity (INU) with N4BiasFieldCorrection (Tustison et al. 2010), distributed with ANTs 2.3.3 (Avants et al. 2008, RRID:SCR_004757), and used as T1w-reference throughout the workflow. The T1w-reference was then skull-stripped with a *Nipype* implementation of the antsBrainExtraction.sh workflow (from ANTs), using OASIS30ANTs as target template. Brain tissue segmentation of cerebrospinal fluid (CSF), white-matter (WM) and gray-matter (GM) was performed on the brain-extracted T1w using fast (FSL 5.0.9, RRID:SCR_002823, Zhang, Brady, and Smith 2001). Volume-based spatial normalization to one standard space (MNI152NLin2009cAsym) was performed through nonlinear registration with antsRegistration (ANTs 2.3.3), using brain-extracted versions of both T1w reference and the T1w template. The following template was selected for spatial normalization: *ICBM 152 Nonlinear Asymmetrical template version 2009c* [Fonov et al. (2009), RRID:SCR_008796; TemplateFlow ID: MNI152NLin2009cAsym].

##### Functional data preprocessing

For each of the 4 BOLD runs found per subject (across all tasks and sessions), the following preprocessing was performed. First, a reference volume and its skull-stripped version were generated by aligning and averaging 1 single-band references (SBRefs). Susceptibility distortion correction (SDC) was omitted. The BOLD reference was then co-registered to the T1w reference using flirt (FSL 5.0.9, Jenkinson and Smith 2001) with the boundary-based registration (Greve and Fischl 2009) cost-function. Co-registration was configured with nine degrees of freedom to account for distortions remaining in the BOLD reference. Head-motion parameters with respect to the BOLD reference (transformation matrices, and six corresponding rotation and translation parameters) are estimated before any spatiotemporal filtering using mcflirt (FSL 5.0.9, Jenkinson et al. 2002). BOLD runs were slice-time corrected using 3dTshift from AFNI 20160207 (Cox and Hyde 1997, RRID:SCR_005927). First, a reference volume and its skull-stripped version were generated using a custom methodology of *fMRIPrep*. The BOLD time-series (including slice-timing correction when applied) were resampled onto their original, native space by applying the transforms to correct for head-motion. These resampled BOLD time-series will be referred to as *preprocessed BOLD in original space*, or just *preprocessed BOLD*. The BOLD time-series were resampled into standard space, generating a *preprocessed BOLD run in MNI152NLin2009cAsym space*. First, a reference volume and its skull-stripped version were generated using a custom methodology of *fMRIPrep*. Several confounding time-series were calculated based on the *preprocessed BOLD*: framewise displacement (FD), DVARS and three region-wise global signals. FD was computed using two formulations following Power (absolute sum of relative motions, Power et al. (2014)) and Jenkinson (relative root mean square displacement between affines, Jenkinson et al. (2002)). FD and DVARS are calculated for each functional run, both using their implementations in *Nipype* (following the definitions by Power et al. 2014). The three global signals are extracted within the CSF, the WM, and the whole-brain masks. Additionally, a set of physiological regressors were extracted to allow for component-based noise correction (*CompCor*, Behzadi et al. 2007). Principal components are estimated after high-pass filtering the *preprocessed BOLD* time-series (using a discrete cosine filter with 128s cut-off) for the two *CompCor* variants: temporal (tCompCor) and anatomical (aCompCor). tCompCor components are then calculated from the top 2% variable voxels within the brain mask. For aCompCor, three probabilistic masks (CSF, WM and combined CSF+WM) are generated in anatomical space. The implementation differs from that of Behzadi et al. in that instead of eroding the masks by 2 pixels on BOLD space, the aCompCor masks are subtracted a mask of pixels that likely contain a volume fraction of GM. This mask is obtained by thresholding the corresponding partial volume map at 0.05, and it ensures components are not extracted from voxels containing a minimal fraction of GM. Finally, these masks are resampled into BOLD space and binarized by thresholding at 0.99 (as in the original implementation). Components are also calculated separately within the WM and CSF masks. For each CompCor decomposition, the *k* components with the largest singular values are retained, such that the retained components’ time series are sufficient to explain 50 percent of variance across the nuisance mask (CSF, WM, combined, or temporal). The remaining components are dropped from consideration. The head-motion estimates calculated in the correction step were also placed within the corresponding confounds file. The confound time series derived from head motion estimates and global signals were expanded with the inclusion of temporal derivatives and quadratic terms for each (Satterthwaite et al. 2013). Frames that exceeded a threshold of 0.5 mm FD or 1.5 standardised DVARS were annotated as motion outliers. All resamplings can be performed with *a single interpolation step* by composing all the pertinent transformations (i.e. head-motion transform matrices, susceptibility distortion correction when available, and co-registrations to anatomical and output spaces). Gridded (volumetric) resamplings were performed using antsApplyTransforms (ANTs), configured with Lanczos interpolation to minimize the smoothing effects of other kernels (Lanczos 1964). Non-gridded (surface) resamplings were performed using mri_vol2surf (FreeSurfer).

Many internal operations of *fMRIPrep* use *Nilearn* 0.6.2 (Abraham et al. 2014, RRID:SCR_001362), mostly within the functional processing workflow. For more details of the pipeline, see <u>the section corresponding to workflows in *fMRIPrep*’s documentation</u>.

##### Copyright Waiver

The above boilerplate text was automatically generated by fMRIPrep with the express intention that users should copy and paste this text into their manuscripts *unchanged*. It is released under the <u>CC0</u> license.

## References

Anderson, R. C. (1978). Schema-Directed Processes in Language Comprehension. In Cognitive Psychology and Instruction (pp. 67–82). Springer US. 10.1007/978-1-4684-2535-2_8

Bailey, H. R., & Zacks, J. M. (2015). Situation Model Updating in Young and Older Adults: Global versus Incremental Mechanisms. Psychology and Aging, 30(2), 232–244. 10.1037/a0039081

Baldassano, C., Chen, J., Zadbood, A., Pillow, J. W., Hasson, U., & Norman, K. A. (2017). Discovering Event Structure in Continuous Narrative Perception and Memory. Neuron, 95(3), 709–721.e5. 10.1016/j.neuron.2017.06.041

Baldwin, D. A., & Kosie, J. E. (2021). How Does the Mind Render Streaming Experience as Events? Topics in Cognitive Science, 13(1), 79–105. 10.1111/tops.12502

Bartlett, F. C. (1932). Remembering: A study in experimental and social psychology. Cambridge university press.

Ben-Yakov, A., & Dudai, Y. (2011). Constructing realistic engrams: Poststimulus activity of hippocampus and dorsal striatum predicts subsequent episodic memory. The Journal of Neuroscience, 31(24), 9032–9042. 10.1523/JNEUROSCI.0702-11.2011

Ben-Yakov, A., & Henson, R. N. (Eds.). (2018). The Hippocampal Film Editor: Sensitivity and Specificity to Event Boundaries in Continuous Experience. The Journal of Neuroscience, 38(47), 10057–10068. 10.1523/JNEUROSCI.0524-18.2018

Bezdek, M. A., Nguyen, T. T., Hall, C. S., Braver, T. S., Bobick, A. F., & Zacks, J. M. (2022). The multi-angle extended three-dimensional activities (META) stimulus set: A tool for studying event cognition. Behavior Research Methods. 10.3758/s13428-022-01980-8

Brunec, I. K., Moscovitch, M., & Barense, M. D. (2018). Boundaries Shape Cognitive Representations of Spaces and Events. Trends in Cognitive Sciences, 22(7), 637–650. 10.1016/j.tics.2018.03.013

Burunat, I., Levitin, D. J., & Toiviainen, P. (2024). Breaking (musical) boundaries by investigating brain dynamics of event segmentation during real-life music-listening. Proceedings of the National Academy of Sciences, 121(36), e2319459121. 10.1073/pnas.2319459121

Chang, C. H. C., Nastase, S. A., & Hasson, U. (2021). Information flow across the cortical timescales hierarchy during narrative construction. 10.1101/2021.12.01.470825

Clark, A. (2013). Whatever next? Predictive brains, situated agents, and the future of cognitive science. Behavioral and Brain Sciences, 36(3), 181–204. 10.1017/S0140525X12000477

Clewett, D., DuBrow, S., & Davachi, L. (2019). Transcending time in the brain: How event memories are constructed from experience. Hippocampus, 29(3), 162–183. 10.1002/hipo.23074

Ding, Y., & Zacks, J. M. (2025). Temporal order memory in naturalistic events is scaffolded by semantic knowledge and hierarchical event structure. 10.31234/osf.io/scaf8_v1

Dosenbach, N. U. F., Koller, J. M., Earl, E. A., Miranda-Dominguez, O., Klein, R. L., Van, A. N., Snyder, A. Z., Nagel, B. J., Nigg, J. T., Nguyen, A. L., Wesevich, V., Greene, D. J., & Fair, D. A. (2017). Real-time motion analytics during brain MRI improve data quality and reduce costs. NeuroImage, 161, 80–93. 10.1016/j.neuroimage.2017.08.025

DuBrow, S., & Davachi, L. (2013). The influence of context boundaries on memory for the sequential order of events. Journal of Experimental Psychology: General, 142(4), 1277–1286. 10.1037/a0034024

DuBrow, S., Rouhani, N., Niv, Y., & Norman, K. A. (2017). Does mental context drift or shift? Current Opinion in Behavioral Sciences, 17, 141–146. 10.1016/j.cobeha.2017.08.003

Etzel, J. A. (2023). Efficient evaluation of the Open QC task fMRI dataset. Frontiers in Neuroimaging.

Feinberg, D. A., Moeller, S., Smith, S. M., Auerbach, E., Ramanna, S., Miller, K. L., Ugurbil, K., & Yacoub, E. (2010). Multiplexed Echo Planar Imaging for Sub-Second Whole Brain FMRI and Fast Diffusion Imaging. PLoS ONE, 5(12).

Friston, K. (2005). A theory of cortical responses. Philosophical Transactions of the Royal Society B: Biological Sciences, 360(1456), 815–836. 10.1098/rstb.2005.1622

Friston, K., FitzGerald, T., Rigoli, F., Schwartenbeck, P., O’Doherty, J., & Pezzulo, G. (2016). Active inference and learning. Neuroscience & Biobehavioral Reviews, 68, 862–879. 10.1016/j.neubiorev.2016.06.022

Gal, Y., & Ghahramani, Z. (2016). Dropout as a Bayesian Approximation: Representing Model Uncertainty in Deep Learning. Proceedings of the 33 Rd International Conference on Machine Learning.

Geerligs, L., Gözükara, D., Oetringer, D., Campbell, K. L., Van Gerven, M., & Güçlü, U. (2022). A partially nested cortical hierarchy of neural states underlies event segmentation in the human brain. eLife, 11, e77430. 10.7554/eLife.77430

Geerligs, L., van Gerven, M., & Güçlü, U. (2021). Detecting neural state transitions underlying event segmentation. NeuroImage, 236, 118085. 10.1016/j.neuroimage.2021.118085

Graesser, A. C., & Nakamura, G. V. (1982). The Impact of a Schema on Comprehension and Memory. In Psychology of Learning and Motivation (Vol. 16, pp. 59–109). Elsevier. 10.1016/S0079-7421(08)60547-2

Gumbsch, C., Adam, M., Elsner, B., Martius, G., & Butz, M. V. (2022). Developing hierarchical anticipations via neural network-based event segmentation. 2022 IEEE International Conference on Development and Learning (ICDL), 1–8. 10.1109/ICDL53763.2022.9962224

Gurney, J. (2017). The Washington University Central Neuroimaging Data Archive.

Hasson, U., Chen, J., & Honey, C. J. (2015). Hierarchical process memory: Memory as an integral component of information processing. Trends in Cognitive Sciences, 19(6), 304–313. 10.1016/j.tics.2015.04.006

Kendall, A., & Gal, Y. (2017). What Uncertainties Do We Need in Bayesian Deep Learning for Computer Vision? Advances in Neural Information Processing Systems.

Knott, A., & Takac, M. (2021). Roles for Event Representations in Sensorimotor Experience, Memory Formation, and Language Processing. Topics in Cognitive Science, 13(1), 187–205. 10.1111/tops.12497

Kuperberg, G. R. (2021). Tea With Milk? A Hierarchical Generative Framework of Sequential Event Comprehension. Topics in Cognitive Science, 13(1), 256–298. 10.1111/tops.12518

Kurby, C. A., & Zacks, J. M. (2018). Preserved neural event segmentation in healthy older adults. Psychology and Aging, 33(2), 232–245. 10.1037/pag0000226

Magliano, J., Trabasso, T., & Graesser, A. (1999). Strategic Processing During Comprehension. Journal of Educational Psychology, 91, 615–629. 10.1037/0022-0663.91.4.615

Nguyen, T. T., Bezdek, M. A., Gershman, S. J., Bobick, A. F., Braver, T. S., & Zacks, J. M. (2024). Modeling human activity comprehension at human scale: Prediction, segmentation, and categorization. PNAS Nexus, 3(10), pgae459. 10.1093/pnasnexus/pgae459

Niv, Y., & Schoenbaum, G. (2008). Dialogues on prediction errors. Trends in Cognitive Sciences, 12(7), 265–272. 10.1016/j.tics.2008.03.006

Radvansky, G. A., & Copeland, D. E. (2006). Walking through doorways causes forgetting: Situation models and experienced space. Memory & Cognition, 34(5), 1150–1156. 10.3758/BF03193261

Radvansky, G. A., Tamplin, A. K., & Krawietz, S. A. (2010). Walking through doorways causes forgetting: Environmental integration. Psychonomic Bulletin & Review, 17(6), 900–904. 10.3758/PBR.17.6.900

Richmond, L. L., & Zacks, J. M. (2017). Constructing experience: Event models from perception to action. Trends in Cognitive Sciences, 21(12), 962–980. 10.1016/j.tics.2017.08.005

Sargent, J. Q., Zacks, J. M., Hambrick, D. Z., Zacks, R. T., Kurby, C. A., Bailey, H. R., Eisenberg, M. L., & Beck, T. M. (2013). Event segmentation ability uniquely predicts event memory. Cognition, 129(2), 241–255. 10.1016/j.cognition.2013.07.002

Schaefer, A., Kong, R., Gordon, E. M., Laumann, T. O., Zuo, X.-N., Holmes, A. J., Eickhoff, S. B., & Yeo, B. T. T. (2018). Local-Global Parcellation of the Human Cerebral Cortex from Intrinsic Functional Connectivity MRI. Cerebral Cortex, 28(9), 3095–3114. 10.1093/cercor/bhx179

Speer, N. K., Zacks, J. M., & Reynolds, J. R. (2007). Human Brain Activity Time-Locked to Narrative Event Boundaries. Psychological Science, 18(5), 449–455. 10.1111/j.1467-9280.2007.01920.x

Su, X., & Swallow, K. M. (2024). People can reliably detect action changes and goal changes during naturalistic perception. Memory & Cognition, 52(5), 1093–1111. 10.3758/s13421-024-01525-8

Swallow, K. M., Zacks, J. M., & Abrams, R. A. (2009). Event boundaries in perception affect memory encoding and updating. Journal of Experimental Psychology: General, 257.

Taylor, A. J., Kim, J. H., & Ress, D. (2018). Characterization of the hemodynamic response function across the majority of human cerebral cortex. NeuroImage, 173, 322–331. 10.1016/j.neuroimage.2018.02.061

Wang, Y. C., & Egner, T. (2022). Switching task sets creates event boundaries in memory. Cognition, 221, 104992. 10.1016/j.cognition.2021.104992

Xu, J., Moeller, S., Auerbach, E. J., Strupp, J., Smith, S. M., Feinberg, D. A., Yacoub, E., & Uğurbil, K. (2013). Evaluation of slice accelerations using multiband echo planar imaging at 3T. NeuroImage, 83, 991–1001. 10.1016/j.neuroimage.2013.07.055

Zacks, J. M. (2010). The brain’s cutting-room floor: Segmentation of narrative cinema. Frontiers in Human Neuroscience, 4. 10.3389/fnhum.2010.00168

Zacks, J. M. (2020). Event perception and memory. Annual Review of Psychology, 71(1), 165–191. 10.1146/annurev-psych-010419-051101

Zacks, J. M., Braver, T. S., Sheridan, M. A., Donaldson, D. I., Snyder, A. Z., Ollinger, J. M., Buckner, R. L., & Raichle, M. E. (2001). Human brain activity time-locked to perceptual event boundaries. Nature Neuroscience, 4(6), 651–655. 10.1038/88486

Zacks, J. M., Speer, N. K., Swallow, K. M., Braver, T. S., & Reynolds, J. R. (2007). Event perception: A mind-brain perspective. Psychological Bulletin, 133(2), 273–293.

## References (from fMRIprep boilerplate)

Abraham, Alexandre, Fabian Pedregosa, Michael Eickenberg, Philippe Gervais, Andreas Mueller, Jean Kossaifi, Alexandre Gramfort, Bertrand Thirion, and Gael Varoquaux. 2014. “Machine Learning for Neuroimaging with Scikit-Learn.” Frontiers in Neuroinformatics 8. 10.3389/fninf.2014.00014.

Avants, B.B., C.L. Epstein, M. Grossman, and J.C. Gee. 2008. “Symmetric Diffeomorphic Image Registration with Cross-Correlation: Evaluating Automated Labeling of Elderly and Neurodegenerative Brain.” Medical Image Analysis 12 (1): 26–41. 10.1016/j.media.2007.06.004.

Behzadi, Yashar, Khaled Restom, Joy Liau, and Thomas T. Liu. 2007. “A Component Based Noise Correction Method (CompCor) for BOLD and Perfusion Based fMRI.” NeuroImage 37 (1): 90–101. 10.1016/j.neuroimage.2007.04.042.

Cox, Robert W., and James S. Hyde. 1997. “Software Tools for Analysis and Visualization of fMRI Data.” NMR in Biomedicine 10 (4-5): 171–78. 10.1002/(SICI)1099-1492(199706/08)10:4/5<171::AID-NBM453>3.0.CO;2-L.

Esteban, Oscar, Ross Blair, Christopher J. Markiewicz, Shoshana L. Berleant, Craig Moodie, Feilong Ma, Ayse Ilkay Isik, et al. 2018. “FMRIPrep.” Software. Zenodo. 10.5281/zenodo.852659.

Esteban, Oscar, Christopher Markiewicz, Ross W Blair, Craig Moodie, Ayse Ilkay Isik, Asier Erramuzpe Aliaga, James Kent, et al. 2018. “fMRIPrep: A Robust Preprocessing Pipeline for Functional MRI.” Nature Methods. 10.1038/s41592-018-0235-4.

Fonov, VS, AC Evans, RC McKinstry, CR Almli, and DL Collins. 2009. “Unbiased Nonlinear Average Age-Appropriate Brain Templates from Birth to Adulthood.” NeuroImage 47, Supplement 1: S102. 10.1016/S1053-8119(09)70884-5.

Gorgolewski, K., C. D. Burns, C. Madison, D. Clark, Y. O. Halchenko, M. L. Waskom, and S. Ghosh. 2011. “Nipype: A Flexible, Lightweight and Extensible Neuroimaging Data Processing Framework in Python.” Frontiers in Neuroinformatics 5: 13. 10.3389/fninf.2011.00013.

Gorgolewski, Krzysztof J., Oscar Esteban, Christopher J. Markiewicz, Erik Ziegler, David Gage Ellis, Michael Philipp Notter, Dorota Jarecka, et al. 2018. “Nipype.” Software. Zenodo. 10.5281/zenodo.596855.

Greve, Douglas N, and Bruce Fischl. 2009. “Accurate and Robust Brain Image Alignment Using Boundary-Based Registration.” NeuroImage 48 (1): 63–72. 10.1016/j.neuroimage.2009.06.060.

Jenkinson, Mark, Peter Bannister, Michael Brady, and Stephen Smith. 2002. “Improved Optimization for the Robust and Accurate Linear Registration and Motion Correction of Brain Images.” NeuroImage 17 (2): 825–41. 10.1006/nimg.2002.1132.

Jenkinson, Mark, and Stephen Smith. 2001. “A Global Optimisation Method for Robust Affine Registration of Brain Images.” Medical Image Analysis 5 (2): 143–56. 10.1016/S1361-8415(01)00036-6.

Lanczos, C. 1964. “Evaluation of Noisy Data.” Journal of the Society for Industrial and Applied Mathematics Series B Numerical Analysis 1 (1): 76–85. 10.1137/0701007.

Power, Jonathan D., Anish Mitra, Timothy O. Laumann, Abraham Z. Snyder, Bradley L. Schlaggar, and Steven E. Petersen. 2014. “Methods to Detect, Characterize, and Remove Motion Artifact in Resting State fMRI.” NeuroImage 84 (Supplement C): 320–41. 10.1016/j.neuroimage.2013.08.048.

Satterthwaite, Theodore D., Mark A. Elliott, Raphael T. Gerraty, Kosha Ruparel, James Loughead, Monica E. Calkins, Simon B. Eickhoff, et al. 2013. “An improved framework for confound regression and filtering for control of motion artifact in the preprocessing of resting-state functional connectivity data.” NeuroImage 64 (1): 240–56. 10.1016/j.neuroimage.2012.08.052.

Tustison, N. J., B. B. Avants, P. A. Cook, Y. Zheng, A. Egan, P. A. Yushkevich, and J. C. Gee. 2010. “N4ITK: Improved N3 Bias Correction.” IEEE Transactions on Medical Imaging 29 (6): 1310–20. 10.1109/TMI.2010.2046908.

Zhang, Y., M. Brady, and S. Smith. 2001. “Segmentation of Brain MR Images Through a Hidden Markov Random Field Model and the Expectation-Maximization Algorithm.” IEEE Transactions on Medical Imaging 20 (1): 45–57. 10.1109/42.906424.

